# Improved computational analysis of ribosome dynamics from 5’P degradome data using fivepeseq

**DOI:** 10.1101/2020.01.22.915421

**Authors:** Lilit Nersisyan, Maria Ropat, Vicent Pelechano

## Abstract

In eukaryotes, 5’-3’ co-translation degradation machinery follows the last translating ribosome providing an *in vivo* footprint of its position. Thus 5’P degradome sequencing, in addition to informing about RNA decay, also provides valuable information regarding ribosome dynamics. Multiple experimental methods have been developed to investigate the mRNA degradome, however computational tools for their reproducible analysis are lacking. Here we present fivepseq: an easy-to-use application for analysis and interactive visualization of 5’P degradome data. This tool performs both metagene and gene specific analysis, and allows to easily investigate codon specific ribosome pauses. To demonstrate its ability to provide new biological information, we investigate gene specific ribosome pauses in *S. cerevisiae* after eIF5A depletion. In addition to identifying pauses at expected codon motifs, we identify multiple genes with strain-specific frameshifts. To show its wide applicability, we investigate more complex 5’P degradome from *A. thaliana* and discover both motif-specific ribosome protection associated with particular developmental stages, as well as generally increased ribosome protection at termination level associated with age. Our work shows how the use of improved analysis tools for the study of 5’P degradome can significantly increase the biological information that can be derived from such datasets and facilitate its reproducible analysis.

**KEY POINTS:**

- Analysis of 5’P degradome data with fivepseq informs about global and gene-specific translational features.
- Frameshifts in translation-related genes in *S. cerevisiae* may be linked to ribosome stalling.
- Ribosome protection at termination and codon motifs are linked to development in *A. Thaliana*.

## INTRODUCTION

The functional status of living cells largely depends on regulation of the pool of translating mRNAs, realized via opposing mechanisms of transcription and RNA decay. In Eukaryotes, general mRNA decay starts by poly(A) tail shortening followed by 5’-3’ or 3’-5’ decay (1). 5’-3’ decay, whereby the exonuclease XRN1 degrades 5’ monophosphorylated (5’P) mRNA intermediates after decapping, is considered to be the main contributing factor in cytoplasmic mRNA turnover (1). Although initially considered independent events, multiple evidence has now demonstrated that translation and mRNA decay are interconnected processes and that co-translational mRNA degradation is a general phenomenon (2–5). The interaction between the translation and decay machinery occurs so close that the positions of 5’P co-translational mRNA degradation intermediates can be used as a proxy for ribosome dynamics, as we and others have shown in yeast (6–8) and plants (4, 9, 10). In particular, XRN1-driven 5’-3’ mRNA degradation is linked to the movement of the last translating ribosome allowing to obtain an *in vivo* footprint of the ribosome position. This interaction has been characterized also at the structural level showing how mRNA is channeled from the ribosome decoding site directly into the active center of the exonuclease (11). Ribosome profiling, a method based on cellular extraction followed by *in vitro* RNA degradation and sequencing, is the current standard to investigate genome-wide ribosome protection (12, 13). However, investigating the 5’P degradome is a very useful complementary approach and allows to obtain a drug-free measurement of *in vivo* ribosome position (omitting *in vitro* RNA degradation) and to focus on those mRNAs that are undergoing co-translational decay (3, 8, 14). Over the years, multiple techniques have been developed to investigate 5’P mRNA degradation intermediates (6–8, 15–18). Methods, such as GMUCT (genome-wide mapping of uncapped and cleaved transcripts) (16, 18) and PARE (parallel analysis of RNA ends) (17) were originally developed to investigate endonucleolytic cleavage mediated by microRNA (miRNA) in plants. Similar approaches have also been used to investigate endonucleolytic cleavage in budding yeast. To investigate the link between the ribosome position and mRNA decay, we developed 5PSeq (6, 7) and more recently, an improved version of this approach, HT-5PSeq (8). Interestingly, the re-analysis of 5’P data originally generated to investigate miRNA mediated endonucleolytic cleavage, demonstrated that general 5’P degradome sequencing informs about ribosome position also in plants (4, 18, 19). This highlights that in addition to optimized experimental protocols for 5’P degradome sequencing (3–5, 9), it is necessary to develop reproducible and simplified computational protocols enabling the systematic study of 5’ P m RNA degradation intermediates in respect to the ribosome position.

However, analysis of 5’P degradome sequencing is not trivial. It is commonly performed using custom scripts optimized for the methodology and the organism of interest (3, 7, 9). Although sufficient to derive useful biological information, this makes it difficult to reproduce and share results. Additionally, it demands a high level of computational competence and complicates the biological interpretation by users with limited bioinformatics experience. Contrary to the case of 5’P degradome sequencing, multiple pipelines have been developed to analyze ribosome profiling data (20–23). Those pipelines focus on identification of translational open reading frames (ORFs), differential expression at translational level and ribosome stalling, with the assumption that ribosome protection fragments are markers of translational activity (12, 20). A primary feature of ribosome profiling pipelines is the ability to take into consideration variable lengths of ribosome protected fragments, which mostly range from 20-22 nt or 28-32 nt in length (depending on factors, such as ribosome conformation and use of translocation inhibitors, *in vitro* nuclease digestion, sequence context, *etc.*) and to determine the ribosome P and A sites. This is usually performed by read stratification by length, followed by alignment of same-length reads relative to all the annotated start sites in the genome. The resulting cumulative counts of read 5’ endpoints show a peak at a certain distance from the start codon, which is regarded as the P-site offset - the distance from read 5’ site to the ribosome P site. This offset is either computed for the most abundant read length or for several read lengths separately (20). Such preprocessing steps for read length selection and adjustment, although important for ribosome profiling, are not relevant for 5’P mRNA degradome datasets, as the latter usually produce longer reads (e.g. more than 50nt (8)), where only the 5’ site is functionally relevant, and do not require 5’ adjustment (6–8). Thus, applying ribosome profiling analysis tools to 5’P degradome sequencing data is complex and would require computational preprocessing to trim the reads to 28-32 nt, limiting the advantages of long-read sequencing and requiring bioinformatics expertise. In addition, there is no simple interactive application that allows for easy exploration of 5’-3’ co-translational degradation profiles, something that is essential to facilitate biological inference from 5’P degradome sequencing data.

Here we present fivepseq, a simple python based standalone command line application that performs comprehensive analyses of 5’P degradome datasets and provides interactive visualization features for data exploration. *Fivepseq* allows for reproducible analysis of 5’P degradome in respect to translational features informing about ribosome protection patterns in respect to ORFs and codons, at genome-wide and gene-specific levels. We demonstrate its applicability by investigating 5’P degradome sequencing in budding yeast and plants. Using this improved approach, we identify gene and codon specific ribosome stalls after depletion of eIF5A in budding yeast. In addition, we identify multiple novel frameshift events associated with strain specific ribosome stalling. Finally, we validate our computational analysis strategy in plants by investigating GMUCT 2.0 data in *Arabidopsis thaliana* developmental stages. In addition to developmental stage- and codon-specific regulation of translation elongation, we report a clear increase of ribosome stalling at termination level in an age dependent manner.

## MATERIAL AND METHODS

### Implementation

Fivepseq is written in python 2.7 and can be used with python 2.7 or 3.x in Unix operating systems. The input for fivepseq are alignment, and genome sequence and annotation files. We used the plastid framework (24) for retrieving 5’P counts in respect to annotated start and stop positions of gene coding sequences (CDS). Protein coding genes (or alternative features specified at input) are filtered based on transcript attributes in the annotation files. The general distribution of 5’P counts is used to determine outliers. The counts are assumed to fall within Poisson distribution with the λ parameter defined as the mean of all the counts greater than 0. Counts that fall below nominal probability 0 defined by python package *stats* are considered as outliers and their values are set to the lowest value among all the outliers. The noise removal options are adjustable. Library size normalization is performed accounting for preprocessed counts in the coding regions, and counts are presented as reads per million (RPM).

To obtain metagene counts, we query 100 nt (adjustable) around start and stop positions and combine the position-wise counts across all the genes. To obtain meta-count vectors for Fast Fourier transformation (FFT), we align the genes at the start or stop positions, then truncate all the genes to the 0.75 percentile of lengths and add stretches of zeros to shorter genes. The resulting equal length vectors are summed up at each position and the resulting meta-vector is used for FFT analysis implemented in the python *numpy* package. The periodicity values are determined by normalizing the number of waves to the meta-vector length, and the absolute values of the corresponding signals are taken as the measure of periodicity strength. A signal-to-noise ratio is computed for the FFT signal by dividing it to the mean of the rest of the signals, the strongest periodicity statistics are stored as output. To obtain frame preference statistics, we sum the counts falling into the first (F0), the second (F1) and the third (F2) nucleotide in each codon of all the genes. For each frame i, the relative preference compared to the other two frames *j* and *k* is computed as the frame protection index with the formula. Significance of frame-preference is estimated by a t-test comparing the counts in a given frame to the distribution of counts in the other frames. The same calculations are also performed for each gene separately. All the statistics are stored in text files, while the global FPI and the maximum p-value for two possible comparisons in each frame are displayed on the reports.

For each codon, we sum the counts at certain positions upstream from its first nucleotide in all the CDS regions across the genome. By default, we store counts 30 nt upstream and 5 nt downstream (adjustable). For each amino acid, we combine the counts for the set of codons it is encoded by. The same computations are performed for all combinations of two/three consecutive codons (or di/tri-peptides), taking the relative positions of 5’P counts from the first nucleotide of the first codon. The di/tri-codons and di/tri-peptides are then filtered to include the top 50 motifs with highest relative counts at positions −14 nt and −11 nt (adjustable) compared to background distribution in the given region (−30 to +5 nt).

We use the python *bokeh* package (v1.0.4) (25) to visualize the counts and statistics obtained above and export to images (*.png* and *.svg*) and HTML formatted report files. We use the interactive *bokeh* tools for zooming and hovering over features under interest and limiting the view to certain samples.

### Preprocessing

The input for *fivepseq* are alignment (*.bam*) files. To ease the experience for the user, we also provide a script for preprocessing of raw (*.fastq*) files and (*.bam*) alignment file generation (https://github.com/lilit-nersisyan/fivepseq/blob/master/preprocess_scripts/fivepseq_preprocess.sh), which performs (i) adapter trimming with *cutadapt* (26) (with the standard Illumina adapter AGATCGGAAGAGCAC and options -- minimum-length 28 -e 0.2 -o 9 --nextseq-trim 20); (ii) extraction of unique molecular identifiers (UMI) with UMI-tools (27) (with the option -- bc-pattern NNNNNNNN); (iii) quality control before and after *fastq* preprocessing with FastQC [https://www.bioinformatics.babraham.ac.uk/projects/fastqc/] and MultiQC (28); (iv) reference index generation and alignment with STAR (29) (with options - - align Ends Type Extend 5p Of Read1 -- outFilterMatchNmin Over Lread 0.9 -- outFilterMultimapNmax 3 --alignIntronMax 2500); (v) selection of primary alignments for multi-mapped reads with Samtools *view* program (30) (with the option -F 0×100) and indexing with Samtools *index* program; (vi) UMI-based removal of PCR duplicates with UMI-tools (27); (vii) generation of read distribution statistics over different features (rRNA, mRNA, tRNA, snoRNA, snRNA, ncRNA) with bedtools *intersect* program (31); and (viii) generation of lightweight 5’P endpoint distribution (*.bedgraph*) files with bedtools *genomecov* program (31) (with options -bg -5 -strand +/−).

### Datasets

The 5PSeq data from *Saccharomyces cerevisiae* with eIF5A depletion (7) are available from the Gene Expression Omnibus (GEO) under accession GSE91064 (replicates GSM2420386, GSM2420387 and GSM2420390 presented in the main figures). 5PSeq data for *S. cerevisiae* 3AT and CHX treatments and randomly fragmented controls (3) are available under accessions GSM1541731, GSM1541711 and GSM1541717. The *Arabidopsis thaliana* GMUCT 2.0 dataset was available from GSE72505 and the PARE dataset from GSE77549 (9). R64-1-1 and TAIR10 assembiles were used for *S. cerevisiae* and *A. thaliana* genomes. The *fastq* files were preprocessed with the *fivepseq_preprocess.sh* script with its default parameters. UMI extraction and deduplication steps were skipped for the GMUCT 2.0 and PARE datasets and adapter trimming was performed with the options - a TGGAATTCTCGGGTGCCAAGG --minimum-length 20. *Fivepseq* reports were generated with *fivepseq* version 1.0b5 with its default options.

### Motif and frameshift analysis

Counts for the tripeptide motifs were taken from the *tripeptide_pauses.txt* files in the *fivepseq* output. The relative counts at position −11 nt over the background were considered to filter motifs with at least 3-fold enrichment. The logo plots for common motifs were generated with the Seq2Logo generator (32). Motif enrichment for frameshift analysis in RNA sequences was performed with the MEME suite (33) under 1-order model of sequences, restricting motif length to 6-7 nt. Variable-nucleotide and out-of-frame motifs were filtered out.

For comparison of gene-specific frame preferences we took statistics from the *frame_counts_TERM.txt* files in the *fivepseq* output for the wild type and the *tif5A1-3* strains. Only genes with at least 50 reads mapping to at least 30 positions were considered. We took the genes where the dominant frame (the one with highest FPI) was the same across all the replicates in each strain. The frame-specific count line-charts were produced in R by taking the *counts_FULL_LENGTH.txt* file as input. The counts along the gene body were averaged over a 90 nt window and then separated into a vector for each frame and plotted as overlaid line charts. We then performed manual selection of genes for which we could observe change in the frame along the gene body consistently among the replicates in each strain. We then plotted combined counts across the replicates for the 13 genes chosen in this manner.

## RESULTS

### *Fivepseq* facilitates reproducible analysis of 5’P degradome data

To address the current limitation of computational methods to analyze and interpret 5’P mRNA degradome sequencing data we developed *fivepseq*, a python based standalone command line application that performs comprehensive analyses of 5’P endpoint distribution and facilitates its biological interpretation. We designed *fivepseq* to generate full reports regarding translational features with a single command (“*fivepseq*”) indicating the mapped reads, and the sequence and annotated features of the genome of interest (Figure 1). Although normally most users will already have reads mapped to their genome of interest, to further facilitate its usability we have also included an auxiliary script (“*fivepseq_preprocess.sh*”) to perform all required preprocessing steps from raw sequencing data (*.fastq*), such as adapter and quality trimming, removal of PCR duplicates and generation of alignment files (*.bam*) (see methods for details). This step also generates lightweight files (*.bedgraph*) to enable visualization of the 5’P endpoints in a genome browser. To facilitate the study of co-translational degradation features, *f* ivepseq automatically masks reads mapping to rRNA, tRNA or non-coding RNA (adjustable upon user input). With this information *fivepseq* generates interactive reports in HTML format (see the report links in the data availability section), providing tools for navigation, zooming, hovering and sample selection, using the *bokeh* package as framework (25). To facilitate data sharing and downstream analysis we have designed *fivepseq* to output also text files with count distribution and statistics, and to generate publication quality vector images.

**Figure 1.**
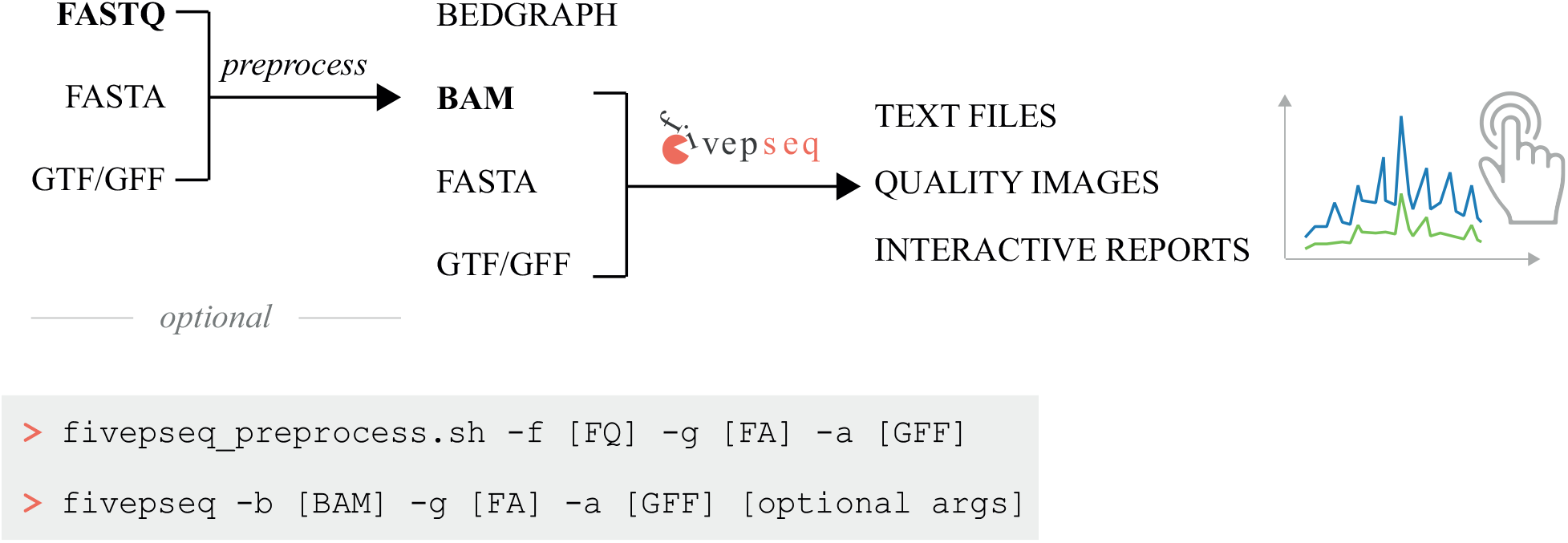
Schematic representation of the fivepseq workflow. The preprocessing steps generate alignment (*.bam*) files from raw reads (*.fastq*) and may optionally be performed with the preprocessing script *fivepseq_preprocess.sh*. It also produces lightweight files (*.bedgraph*) for visualization in genome browsers. The main fivepseq workflow takes alignment and genome sequence and annotation files as input and generates interactive reports and publication quality image files describing the translational features, as well as stores counts and statistics in text files suitable for downstream analysis. Type to enter text

In brief, *fivepseq* performs the following analysis. First, we analyze the mapped reads to obtain 5’P count vectors in respect to annotated start and stop positions of gene coding sequences (CDS) using *plastid* as a framework and sum the counts at each position across genes to obtain metagene information (Figure 2A) (24). As *fivepseq* has been designed to investigate *in vivo* 5’-3’ footprints generated by the cellular degradation machinery, we decided not to apply any correction to the 5’ end position, unlike many ribosome profiling approaches (12, 20). We reasoned that any observed variation will be caused by *in vivo* physiological variations (not due to differential *in vitro* RNA digestion) and decided to provide the user with real protection patterns to facilitate the biological interpretation with no prior assumptions regarding fragment position relative to the ribosome. To avoid the effect of outliers, we perform data cleaning and noise reduction steps, down-scaling extremely high 5’P counts, which adds to robustness of *fivepseq*.

**Figure 2.**
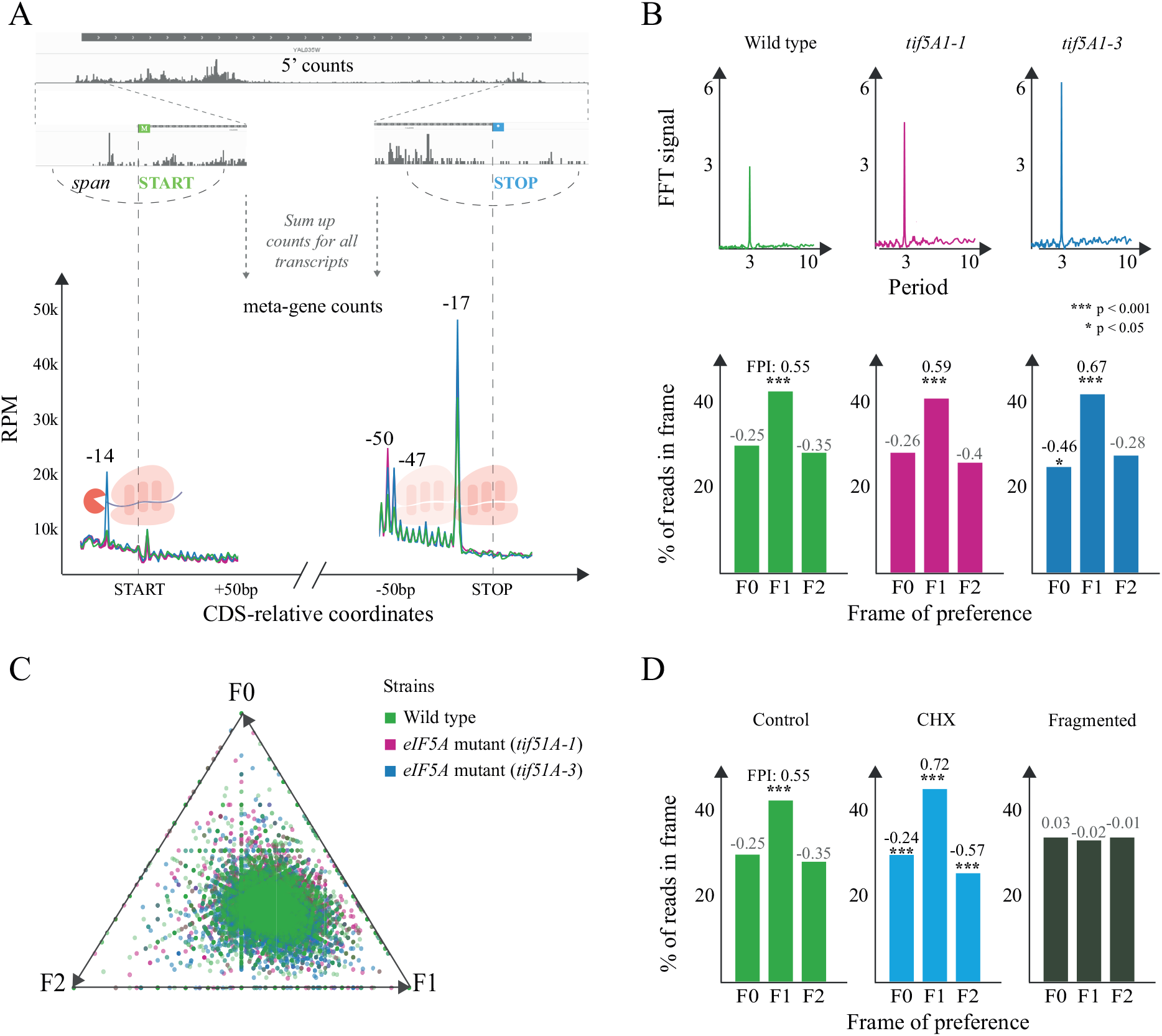
General 5’P degradome distribution in budding yeast. **A.** For each transcript, fivepseq queries a region around the CDS start and stop codons and gets a vector containing 5’P count information. These vectors are summed up to derive a metagene coverage around the start and stop codons. Library size normalized counts are presented as reads per million (RPM) considering only those reads mapping in the coding regions. **B.** Top: Fast Fourier transform (FFT) analysis showing the strength (y-axis) and periodicity (x-axis) values. Bottom: Histograms displaying the global frame preference for the first (F0), the second (F1) and the third (F2) nucleotide of each codon. The frame protection index (FPI) shows the strength of preference for each frame along with a t-test p-value comparing the counts to the other two frames. **C.** Frame preference values for each gene converted to 2D space in a triangle plot, where each point is a gene and its distance from the triangle vertex is inversely proportional to respective frame preference. **D.** Genome wide frame preferences for a positive control, a CHX treated sample and a randomly fragmented negative control (3).

As ribosomes move along the mRNAs one codon at a time, we also analyze the presence of 3-nt periodicity and frame preference in the 5’P degradome data. We sum the counts falling into the first (F0), the second (F1) or the third (F2) nucleotide of each codon in each gene and represent it as a histogram. To quantify variations in the observed frame protection patterns we also provide statistics for frame protection - the frame protection index (FPI, see methods) and a t-test p-value comparing counts in each frame to the other two. In addition to global profiles, 5’P degradome sequencing can also provide gene-specific information. To achieve this, we transform the gene-specific frame counts into 2D coordinates and visualize them with triangle plots, where each point is a gene and each triangle vertex a frame (Figure 2B). The higher the counts in frames F0, F1 or F2, the closer is the point to the respective triangle vertex. To provide a more sensitive measure of presence of periodic count patterns without restricting to 3-nt, we also perform a Fast Fourier transformation based (FFT) analysis. As anticipated, the FFT analysis shows a clear periodicity peaking at 3-nt, as expected from the protection patterns of a ribosome moving one codon at a time (Figure 2B).

In addition to general ribosome protection patterns associated with translation initiation and termination, it is also important to identify codon-specific protection patterns arising in translation elongation. Context-dependent ribosome protection (or stalling) can inform, for example, about differential velocity for incorporation of certain amino acids (3) or even interactions between the ribosome exit tunnel and particular peptide sequences (7). This effect can be even more drastic when targeting amino acid or tRNA metabolism (3, 34). To aid in those analyses, we compute the metagene profiles for 5’P endpoints positions at a certain distance from the first nucleotide of each codon or amino acid (see below Figure 3A). We generate interactive line charts for each individual codon (or amino acid) where samples are overlaid and specific samples may be interactively highlighted or hidden. We also summarize that information in a heatmap to facilitate the comparison across codons within a sample (Figure 3A). To ease comparison of translational features between samples, we also generate differential heatmaps for each sample pair, where the normalized difference between counts at every position from each codon is displayed (Figure 3B,C). As certain combinations of codons or amino acids can also affect ribosome dynamics (35, 36), we additionally analyzed the ribosome protection associated with the presence of two and three codon combinations (and di/tri-peptide) (Figure 3D-E).

**Figure 3.**
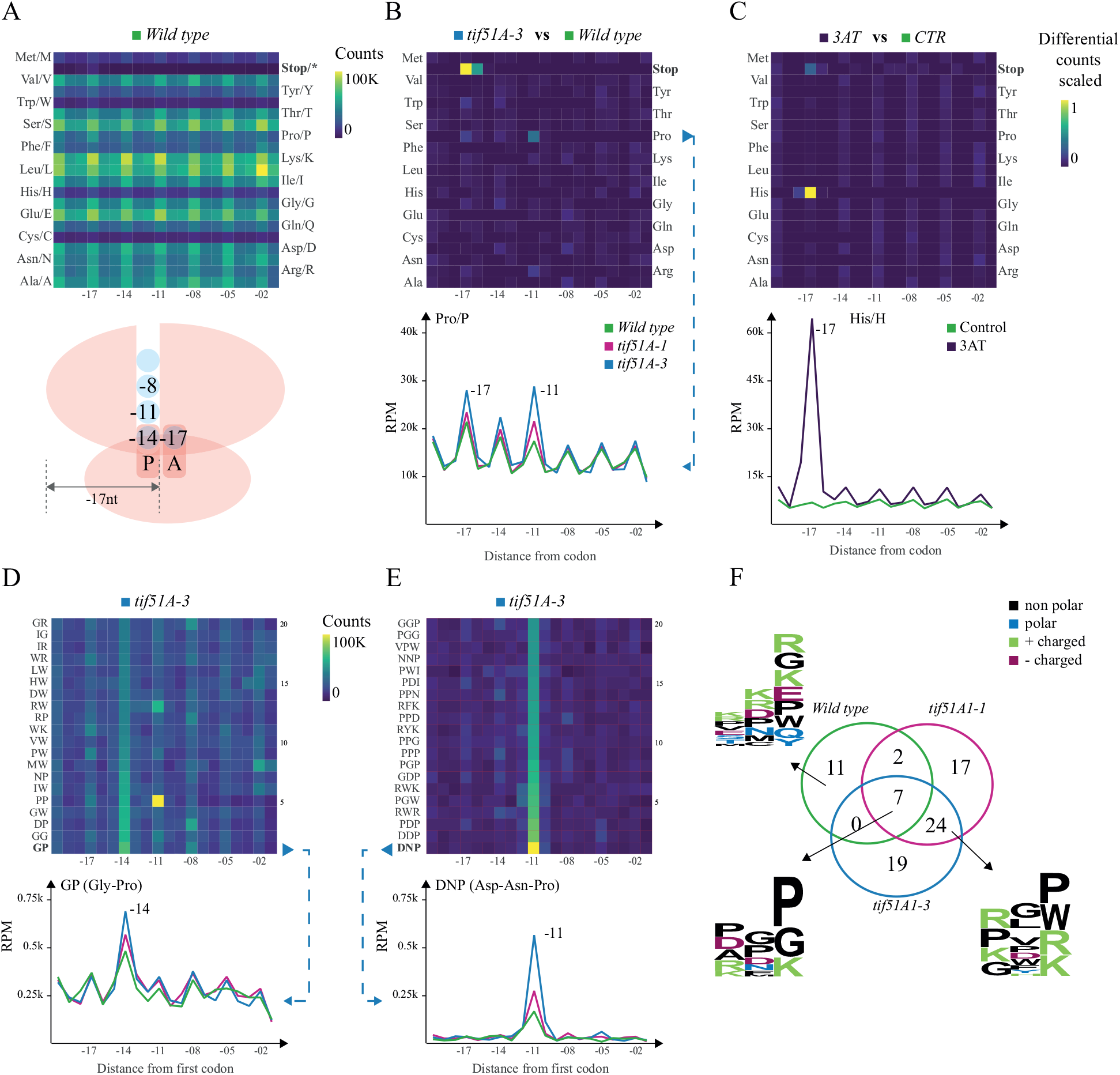
Codon and codon-motif specific ribosome protection patterns in budding yeast. **A.** Heatmap of counts located at a certain distance from each amino acid in the wild type. The ribosome scheme depicts positions of 5’P endpoints relative to the ribosome. **B.** A differential heatmap showing the difference in scaled 5’P counts between the *tif5A1-3 eIF5A* mutant and the wild type for all the amino acids, and a line chart highlighting the difference between both mutants and the wild type for proline (Pro/P). **C.** A differential heatmap of codon-specific 5’P counts between 3AT treated and control yeast cells highlighting the stalling at −17 nt from histidine (His/H) and the corresponding line chart. **D.** The top 20 dipeptides showing relatively high counts at −14 nt in the *tif5A1-3 eIF5A* mutant and the strongest pausing caused by GP (Gly-Pro) dipeptide, comparing the two eIF5A mutants and the wild type. **E.** The top 20 tripeptide motifs showing the strongest −11 nt pausing in the *tif5A1-3 eIF5A* mutant and a line chart comparing the two mutants and the wild type strains for the strongest pause associated tripeptide DNP (Asp-Asn-Pro). **F.** A venn diagram showing the overlap of tripeptides with at least 3-fold increase in 5’P counts at position −11 nt compared to the background, in the three yeast strains. The logo plots present the motifs specific to the wild type, or the three strains or to the two *eIF5A* mutants.

Finally, as in some cases particular groups of genes display differential translational features, we allow for specification of particular gene sets for detailed analysis using a simple text input. This allows for easy comparison of ribosome protection signatures across gene sets and samples.

### Global and gene-specific ribosome protection patterns upon eIF5A depletion

Having developed the *fivepseq* framework, we decided to demonstrate its utility by reanalyzing our recent 5PSeq data targeting the translation elongation factor eIF5A in *S. cerevisiae* (7). eIF5A is thought to act in translation elongation by binding the E-site and promoting peptide bond formation (37). In addition, the lack of eIF5A is known to increase ribosome stalling at proline and stop codons (37–39). We analyzed 5PSeq data obtained from thermosensitive yeast strains carrying one (*tif1A-1*) or two (*tif5A1-3*) point mutations in the *eIF5A* gene (7). As expected, we clearly see a 5’P peak 17 nt upstream from the stop codon, indicating RNA decay fragments protected by ribosomes stalled at termination (Figure 2A). This protection at termination was enhanced for both mutants and led also to an increased protection 47 nt and 50 nt upstream from the stop codon, indicating disomes stalled at the stop codon. We also detected a clear protection pattern 14 nt upstream from the start codon for the *tif5A1-3* mutant that corresponds to ribosome stalling in the P site at the start codon, in agreement with our previous finding (7). In addition to start and stop peaks, we also investigated the variation of 3-nt periodicity across strains, as it can provide information regarding translation elongation speed and co-translational mRNA degradation sensitivity (3). To expand our prior work, we quantitatively assess the presence and compare the strength of genome-wide 3-nt periodicity applying Fast Fourier transformation (FFT) analysis. As seen in Figure 2B, the presence of 3-nt periodicity is apparent in all the three strains and increases in the *eIF5A* mutants. Additionally, we can easily see that the 5’ counts are accumulated in the second nucleotide of each codon (F1), with counts significantly greater than those in frames F0 and F2, and this effect was more apparent in the *eIF5A* mutants (F1 frame protection index (FPI): 0.59 and 0.67 *versus* 0.55 in the wild type). This frame preference observed at metagene level is not driven only by a few genes, as we can see that gene-specific frame protection is generally skewed toward F1 (Figure 2C). To demonstrate the general applicability of these analyses, we also analyzed 5’P degradome after stalling ribosomes with cycloheximide and using randomly fragmented RNA as a negative control, where no apparent 3-nt periodicity was observed (Figure 2D).

### Codon-specific ribosome protection patterns upon eIF5A depletion

The role of eIF5A in translation elongation was initially restricted to its ability to contribute to releasing ribosome stalls at polyproline motifs (39). However, recent data shows that this role is not limited to polyproline motifs, but also combinations of other amino acids, including proline, glycine and charged amino acids (7, 38). To investigate elongation stalls in more detail we first analyzed codon and amino acid specific ribosome protection. To facilitate the comparison of protection patterns for the same codon between different conditions, *fivepseq* generates differential heatmaps, where only the difference of scaled counts between each pair of conditions is shown. For example, the differential heatmap between the *tif5A1-3 eIF5A* mutant (which displays a stronger phenotype compared to *tif5A1-1*) and the wild type, clearly reveals not only increased protection at termination, but also at −17 nt and −14 nt from proline (Figure 3B). Given the side chain structure of proline, it induces ribosome stalls during peptide bond formation, which normally get alleviated by eIF5A (40), while the pause at −11 nt could be explained by differential interaction with the ribosome exit tunnel (7). Similar changes in protection were also observed for glycine at positions −14, for arginine at −11, *etc.* These pauses suggest alleviation of ribosome stalling by eIF5A when the amino acid has already been incorporated into the peptide chain and interacts with the exit tunnel. To demonstrate the robustness of this analysis, we also analyzed ribosome protection after 3AT treatment (3), which inhibits the histidine biosynthesis pathway and induces drastic ribosome stalls at histidine codons (Figure 3C).

In addition to single amino acids, specific arrangement of consecutive amino acids can also lead to ribosome stalls, as the neighboring amino acids affect peptide bond formation or modulate interactions in the ribosome exit tunnel (35). In other cases, specific codon motifs can change the mRNA conformation, interfering with the decoding process (36). To facilitate the exploration of this phenomenon, we generate an automatic report investigating two or three codon (or amino acids) combinations able to produce ribosome stalls. This analysis shows for example, that the dipeptide GP (glycine-proline) induces pausing in the *tif5A1-3 eIF5A* mutant (Figure 3D). More pauses are observed at tripeptide motifs both in the mutants and in the wild type. For example, DNP (Asp-Asn-Pro) shows the strongest stall in the *tif5A1-3* mutant (Figure 3E). Notably, the tripeptides were enriched in prolines in the E and A sites, but other amino acids, such as aspartic acid and arginine were also enriched in these positions. To further expand our analysis and for more systematic characterization of the tripeptides, we focused on those motifs for which more than 3-fold increase in −11 counts was observed. Seven motifs were common in all the samples, while the majority were specific to the mutants. Interestingly, there were also motifs that showed 3-fold increase only in the wild type (Figure 3F, Supplementary Figure S1). Motif analysis showed enrichment of proline, glycine and charged amino acids arginine and lysine in the E and A sites in the *eIF5A* mutants, while in the wild type strain lysine and arginine were frequently observed in the E, P and A sites, and glycine and glutamic acid mainly in the A site (7).

### Contribution of ribosome stalling and eIF5A depletion to ribosome frameshifts

Having confirmed our ability to identify and expand known eIF5A biology, we decided to use fivepseq to investigate gene specific regulation of ribosome protection patterns. We wondered whether the changes in frame protection patterns observed at global level (Figure 2B-C) could also be detected at single-gene resolution. Taking the gene-specific frame preference statistics output of *fivepseq,* we compared frame preference patterns between the wild type and the *tif5A1-3 eIF5A* mutant strains, restricting ourselves to high coverage genes (more than 50 reads distributed along at least 30 different nucleotides). In the wild type strain, we identified 32 genes with 5’P counts predominantly in the frame F0, and 13 genes in the frame F2. Interestingly, those same genes had the expected canonical preference for the frame F1 in the *tif5A1-3* mutant (Supplementary Table S1). In *tif5A1-3*, we identified fewer genes with preference for F0 (6 genes) and F2 (6 genes), while these same genes had F1 preference in the wild type (Supplementary Table S1). To understand if the observed changes in frame preference could partially be explained by ribosome frameshifts, we explored the line charts representing frame-specific counts along the bodies of these genes and manually picked those that consistently changed the frame of preference in the same position in all the replicates for each strain, resulting in 13 genes with likely ribosomal frameshifts (RF) in the wild type in the directions −1 (7 genes, Figure 4A) and +1 (6 genes, Figure 4B) (Supplementary Table S2).

**Figure 4.**
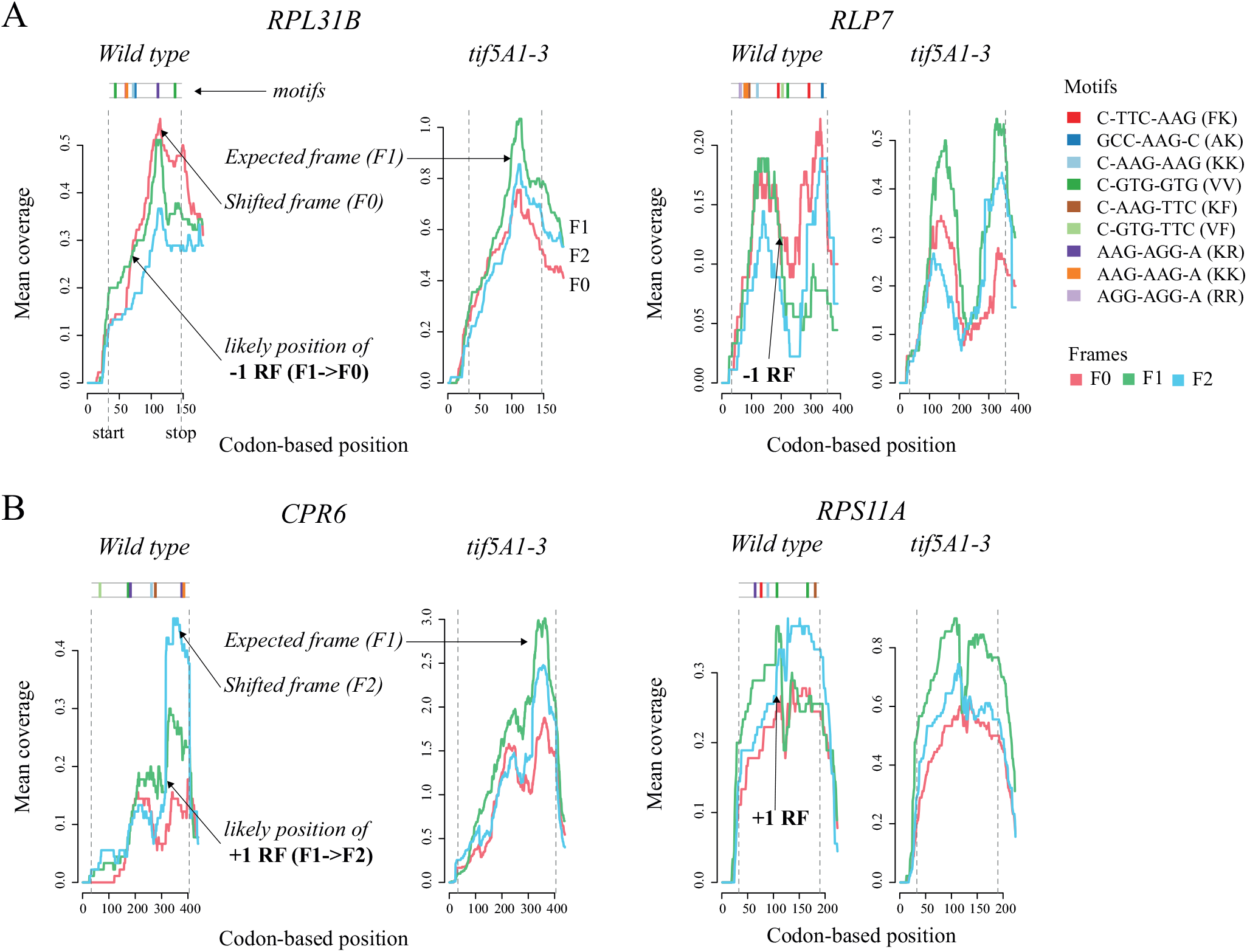
Changes in frame preference patterns in the wild type versus the eIF5A mutant strains. **A.** Two (out of 7) genes, where ribosomal frameshifts (RF) in the −1 direction (from F1 to F0) could be consistently detected across the replicates. The line charts show frame-specific counts (each frame highlighted by a color) averaged over a 30-codon window along the gene body. In the *tif5A1-3* strain the expected (F1) frame is always dominating. In the wild type, starting from some point, the frame F0 becomes predominant, indicating a likely frameshift (−1 RF) at this position. Top: presence of known “slippery” or most enriched 7-nt motifs along the gene body. **B.** Two (out of 6) genes, where ribosomal frameshifts (RF) in the +1 direction (from F1 to F2) could be consistently detected.

As the orthologues or family members of some of these genes were previously reported to experience ribosome frameshifts (*e.g. RPS11A* in *S. pombe,* see Supplementary Table S2), we decided to check for the presence of known “slippery” sequences that commonly promote ribosomal frameshifts. First, we checked for 6 and 7-nucleotide motifs in the RNA sequences, and found enrichment for Lysine and Arginine, in particular for the di-codon AAG-AAG (p-value > 10e-4). Interestingly, di- and tri- peptide motifs enriched in Lys, Arg and Glu showed preferential stalling specifically in the wild type strains, with drastic stalling at poly-lysines (Figure 3F, Figure S1), suggesting possible lack of these amino acids or respective t-RNAs. The observed ribosome stalling in the wild type strain for those genes would lead to ribosome queuing and potentially facilitate frameshifting around “slippery” sequences, as has recently been proposed in bacteria (41). Indeed, 8 out of the13 explored genes had slippery sites (C-TTC-AAG and GCC-AAG-C) that are commonly inducing frameshifts at “hungry” lysine codons (42). The fact that stalling at poly-lysines is preferentially occurring in the wild type (and not the *tif5A1-3* mutant (7)) suggests that the general decrease in relative ribosome load in those genes in the *eIF5A* mutant strain (7) reduces frequencies of ribosome collisions and suppresses frameshifts. This observation might provide functionally relevance to the environmentally regulated stall at SKE motifs (Ser-Lys-Glu) that we described previously in the wild type strains (7). Additionally, the observed enrichment for Arg-Arg sequence AAG-AAG is also known to induce frameshifting in *E. coli*, when the cognate t-RNAs are sparse (42, 43) (Figure 4). Interestingly, all, except for one, of the identified 13 genes code for ribosomal proteins or regulate translation (Supplementary Table S2), suggesting a possible regulatory role (44). Contrary to our expectations, even if the *eIF5A* depletion leads to massive ribosome stalling at polyproline sequence, we were only able to find a few examples where frameshift was induced by *eIF5A* depletion (Supplementary Table S1). 5’P degradome reveals codon-specific ribosome protection patterns in plants

Having demonstrated the ability of *fivepseq* to identify codon- and gene-specific ribosome protection patterns in budding yeast, we decided to apply it to investigate 5’P degradome in *A. thaliana*, an organism of higher complexity. In plants, the cytoplasmic exonuclease XRN4 (orthologue of the yeast XRN1) is responsible for the 5’-3’ co-translational mRNA decay and can also be used to investigate ribosome protection (4, 9). For this purpose, we decided to use a rich GMUCT 2.0 dataset of developmental transitions in *A. thaliana* (Poethig, Meyers, Willmann and McCormick, GSE72505). This dataset was originally generated to study miRNA regulation, and thus was never used to investigate ribosome dynamics. In addition to this, we also explored the PARE data from *A. thaliana* with *fivepseq* (9). However, the use of MmeI during library generation introduces an additional bias that complicates the investigation of codon-specific patterns in these datasets (Figure S2).

We observed clear accumulation of 5’P endpoints 16 nt upstream from the stop codons of all protein coding genes indicating ribosome protection at termination (Figure 5A) and a clear 3-nt periodicity (4, 9). Interestingly, the protection at codons different from the stop is extended by an additional nucleotide (17 nt, as observed in yeast) suggesting that ribosomes stalled at termination level in *A. thaliana* protect a region 1 nt shorter than in budding yeast. Globally, we observed a protection preference for the second nucleotide of each codon (F1) (Figure 5B). The 3-nt patterns were subtler compared to those observed in budding yeast, which may be attributed to more complex biology in plants, longer mRNA half-life (as we hypothesized for *S. pombe* (3, 8)) or differences in 5’P degradome library preparation. Interestingly, we observed a clear regulation of 5’P accumulation associated with the stop codon across developmental stages, which suggests an increase of ribosome termination stall with age (from day 6 to day 33 of growth). This effect was not dependent on genotype (early flowering, flc-3, or late flowering, FLC) or the tissue under study (cotyledons from day 6, apices from days 9 and 11, and leaves from days 14, 23, 32 and 33) (Figure 5A). Increased termination stall during *A. thaliana* development is in line with previously observed reduction in polysome associated mRNAs (45), and the increased ribosome termination stalls that we previously described in budding yeast during limited nutrient conditions or stationary phase (3, 8). Interestingly, apices showed subtle preference for 5’P counts 17nt upstream from the stop codon, as opposed to the general −16 nt preference in the other samples (Figure 5A, Figure S3E). Contrary to previous findings suggesting differential protection for TAA and TAG stop codons (16 and 17 nt respectively) (4), we observe similar preference for all stop codons (TGA, TAG and TAA) (Figure S3E). The coverage surrounding the start codon was relatively low and thus limited our ability to study ribosome protection patterns associated with translation initiation (Figure 5A). This is likely the result of the poly(A) selection strategy used to generate GMUCT 2.0 libraries that biases read coverage towards 3’ ends the genes (4, 6, 8).

**Figure 5.**
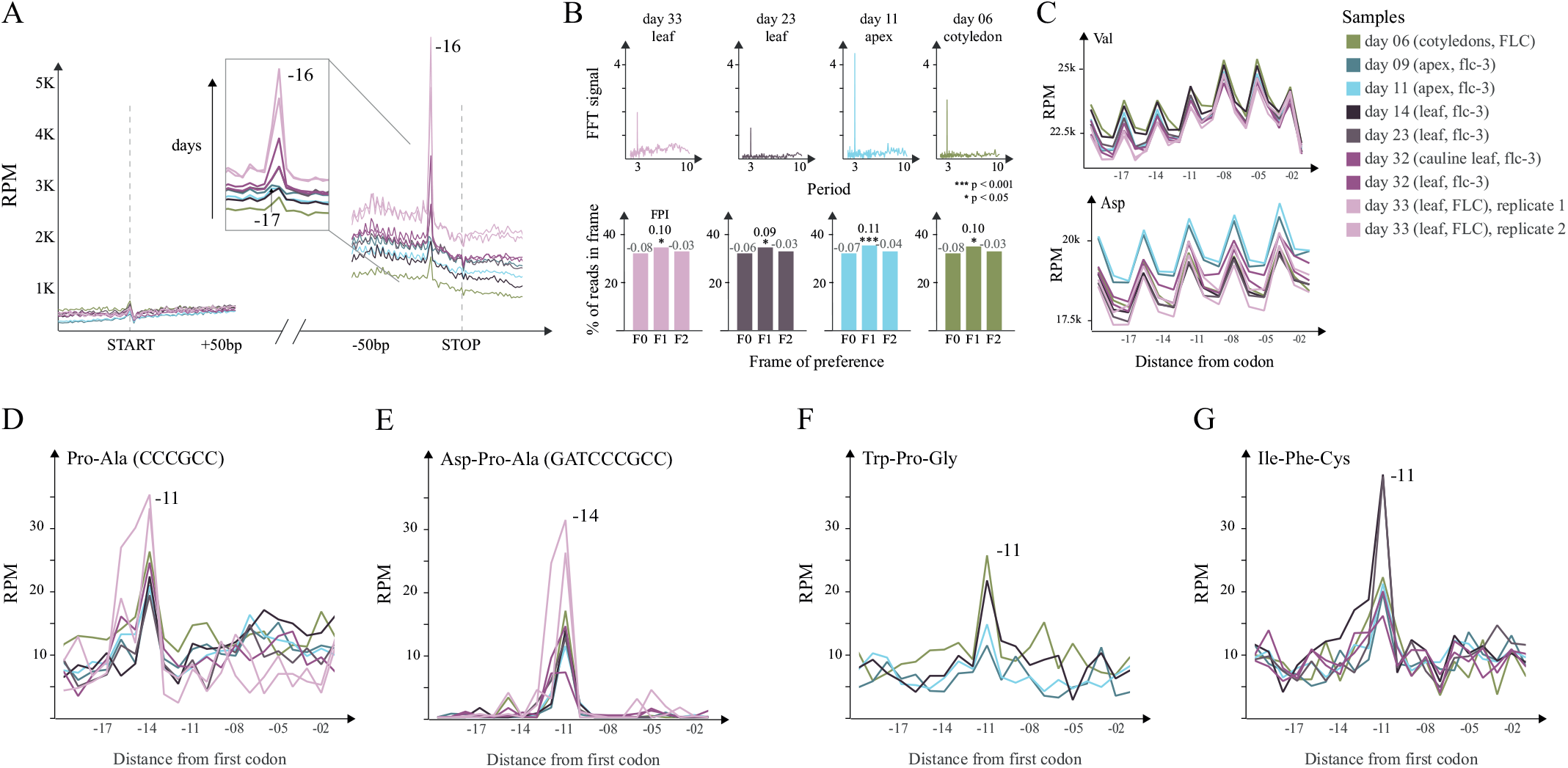
Ribosome protection patterns in Arabidopsis thaliana. **A.** Metagene level protection patterns at start and stop. The stop peak at −17nt is highlighted to underline the increase in protection associated with growth. **B.** Top: Fast Fourier transform (FFT) analysis showing the strength (y-axis) and periodicity (x-axis) values. Bottom: Histograms displaying the global frame preference for the first (F0), the second (F1) and the third (F2) nucleotide of each codon. The frame protection index (FPI) shows the strength of preference for each frame along with a t-test p-value comparing the counts to the other two frames. **C.** Top: Uniform protection patterns at valine across the samples. Bottom: Differential usage of aspartic acid in apices compared to leaves. **D.** Pro-Ala codon pair leading to increased protection at −14 nt in all the samples. **E.** Asp-Pro-Ala tri-codon motif leading to increased protection at −11 nt in all the samples. **F.** The tripeptide Trp-Pro-Gly leading to increased pausing at −11nt in younger samples (days 6, 9, 11 and 14). **G.** The Ile-Phe-Cys tripeptide causes ribosome pausing at −11nt in all the samples except for those grown until day 33.

Having investigated differential co-translational decay associated with translation termination, we moved on to explore differential ribosome protection during translation elongation. Notably, for some amino acids, such as valine, abundances of upstream 5’P reads and protection patterns were similar across all the samples. Whereas for others, such as aspartic acid, a difference in 5’P read abundances was present between leaves and apices (Figure 5C). This suggests differential codon composition on mRNAs expressed at different stages as previously shown (46). At single amino acid level, proline did not induce noticeable ribosome stalling. However, the motifs of two or three codons/ amino acids associated with ribosome pausing, showed enrichment for proline, arginine and glycine, similar to what was observed in yeast (Figure 5D-E, Figure S3A-D). We confirmed accumulation of 5’P reads 14 nt upstream of known inhibitory codon pairs, such as Arg-Arg and Arg-Pro (Figure S3A-C). In addition, we found the Pro-Ala (CCC-GCC) and Asp-Pro-Ala (GAT-CCC-GCC) motifs to be strongly associated with ribosome pausing specifically in *A. thaliana* (Figure 5D-E). This pausing was observed in all the samples and was codon-dependent. We also identified particular motifs where pausing was associated with age. Specifically, the tripeptides Trp-Pro-Gly (WPG) and Ile-Phe-Cys (IFC) (Figure 5F,G), and the dicodon Gln-Leu (CAA-CTG) (Figure S3D) showed increased stalling in earlier developmental stages.

## DISCUSSION

Here we have presented the development of a reproducible computational pipeline for 5’P degradome analysis. *Fivepseq,* performs comprehensive analyses of 5’P degradome data in respect to translational features in a single command line and allows for easy exploration of global and gene-specific translational frame preference and codon-specific ribosome protection patterns. To facilitate its use by scientists with limited bioinformatic expertise, we have implemented interactive features in the visualization reports, which allow for smooth exploratory analysis, such as zooming and hovering over features of interest or easy comparison between samples. In addition, we have also implemented a rich variety of text output files describing count distributions and statistics that can be used by more experienced computational biologists for advanced downstream analysis. *Fivepseq* requires minimal data preprocessing, and also provides a single command line option to convert raw reads to alignment files before main analysis.

To demonstrate robustness and applicability of *fivepseq*, we first re-analyzed previous data from our group focused on the 5’P degradome sequencing in budding yeast after eIF5A depletion (7). We could identify increased ribosome protection at termination level and codon specific ribosome pauses such as DNP (Asp-Asn-Pro) and PPP (Pro-Pro-Pro) associated with eIF5A depletion (7, 38). In addition to known eIF5A biology, our new single gene analysis is able to identify novel features from the original dataset. We identified 57 genes with anomalous frame protection patterns both in the wild-type and the mutant strains. Analysis of those data revealed evidence for frameshifting in 13 genes in the wild type strain associated with the ribosome and translational process. Interestingly the observed frameshifts were associated with combinatorial presence of arginine and lysine rich “slippery” sequences and KKK, RKK motifs that in the wild type strain may induce ribosome pausing (42, 43). This result is in agreement with recent reports showing how ribosome stalling can facilitate frame-shifting events (41). Particularly interesting is the case of the gene *TMA46* that is associated with the resolution of arginine and lysine stalls (14) and for which we also identify a stall dependent frameshift. Suggesting thus a potential cross regulation between both processes. However, it is important to note that 5’P degradome analysis focuses on those molecules undergoing co-translational degradation that represent a subset of the mRNA molecules undergoing active translation. More detailed molecular analysis will be required to confirm both the production of frameshifted proteins and their potential functional role.

Finally, we demonstrate the flexibility of our computational pipeline by analyzing *A. thaliana* GMUCT 2.0 degradome data, originally developed to investigate miRNA regulation during development. In addition to previously described general ribosome protection features (4, 9), we found both similarities and differences in respect to our work in budding yeast. Even though *in vivo* 5’P codon specific ribosome footprints have a similar size between yeast and *A. thaliana*, in the latter ribosome protection at translation termination level is one nucleotide shorter (16 nt in plants in comparison to the 17 nt observed in yeast). Interestingly, we found that increased ribosome protection at translation termination level increases with developmental stage. This phenomenon resembles our observation that in budding yeast limited nutrition or stationary phase also leads to an accumulation of ribosomes stalled at termination level (3, 8). All this is in agreement with an expected decrease in overall translation with age and suggests that translation termination is often regulated (45). In addition to this general regulation, we also identified codon-specific ribosome pausing associated with specific developmental stages.

In summary our work shows how the development of improved computational tools for the analysis of 5’P degradome datasets is critical to derive novel biological insights regarding the crosstalk between translation and mRNA decay. We expect that fivepseq will facilitate the analysis of 5’P degradome sequencing data across multiple organisms and support its reproducible investigation.

## AVAILABILITY

*Fivepseq* is available from http://pelechanolab.com/software/fivepseq, under BSD 3-Clause License. The source code is deposited at https://github.com/lilit-nersisyan/fivepseq and a detailed user manual is available at https://fivepseq.readthedocs.io/en/latest/.

Fivepseq reports generated for this study are available under DOI: https://doi.org/10.17044/scilifelab.12662153

## ACCESSION NUMBERS

Analyzed *Saccharomyces cerevisiae* data are available from GEO under accession codes GSE91064, GSM1541731, GSM1541711 and GSM1541717. *Arabidopsis thaliana* datasets are available under accessions GSE72505 and GSE77549.

## ACKNOWLEDGEMENT

We thank PelechanoLab members for discussions and alpha testing, Arsen Arakelyan (BIG NAS RA), Hrant Khachatrian, Karen Hambardzumyan (YerevaNN, Armenia) and Tigran Shahverdyan for tech-support and math-advice. Computational resources were provided by SNIC through Uppsala Multidisciplinary Center for Advanced Computational Science (UPPMAX).

