## Supplementary material for "Improved computational analysis of ribosome dynamics from 5’P degradome data using fivepeseq"

### SUPPLEMENTARY DATA

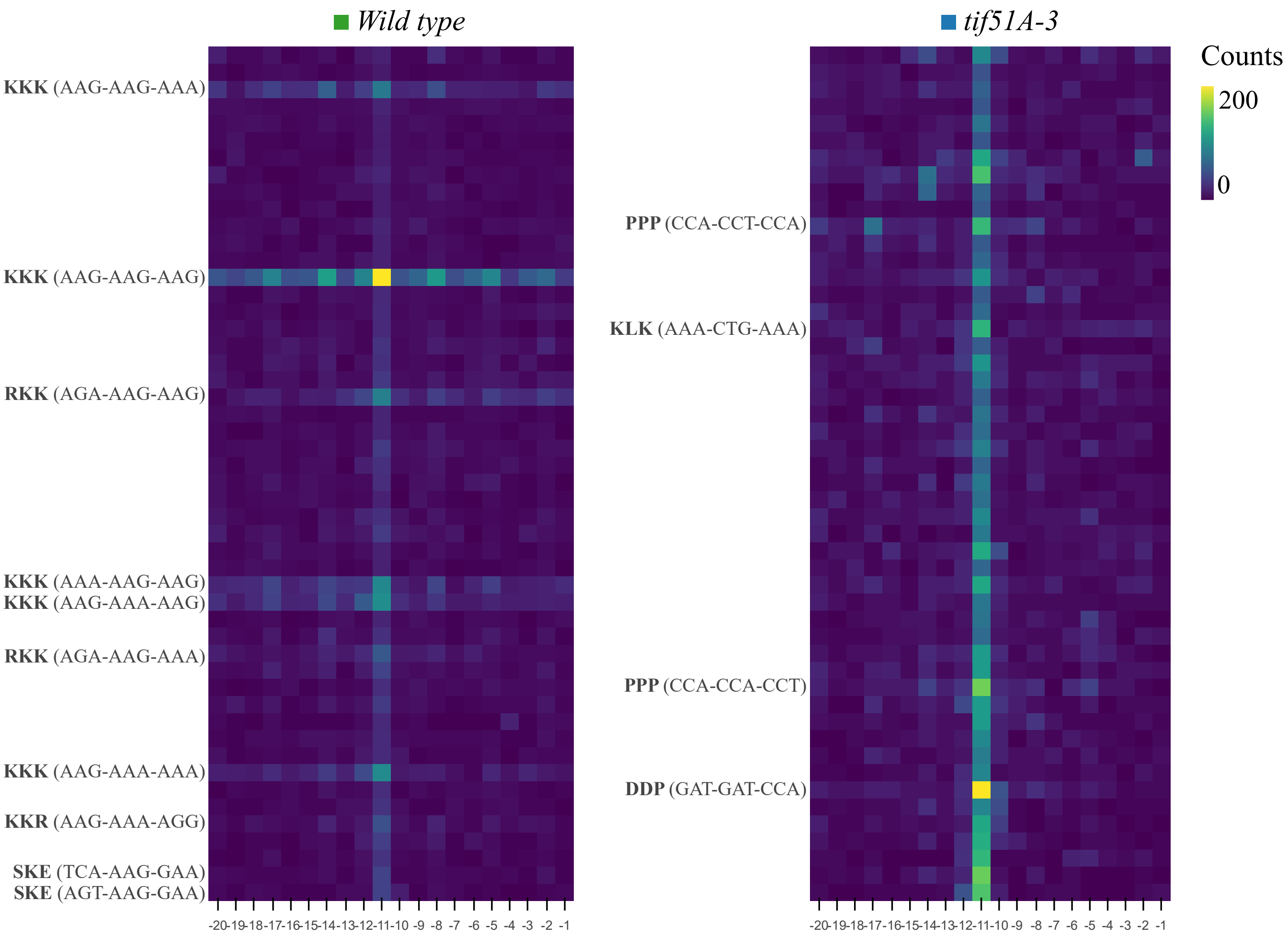


**Figure S1. Tri-codon relative counts in the wild type and the *tif5A1-3* strains.** Heatmaps of counts located at a certain distance from the first codon of top 50 tri-codon motifs sorted by relative counts at position -11 nt over the background.

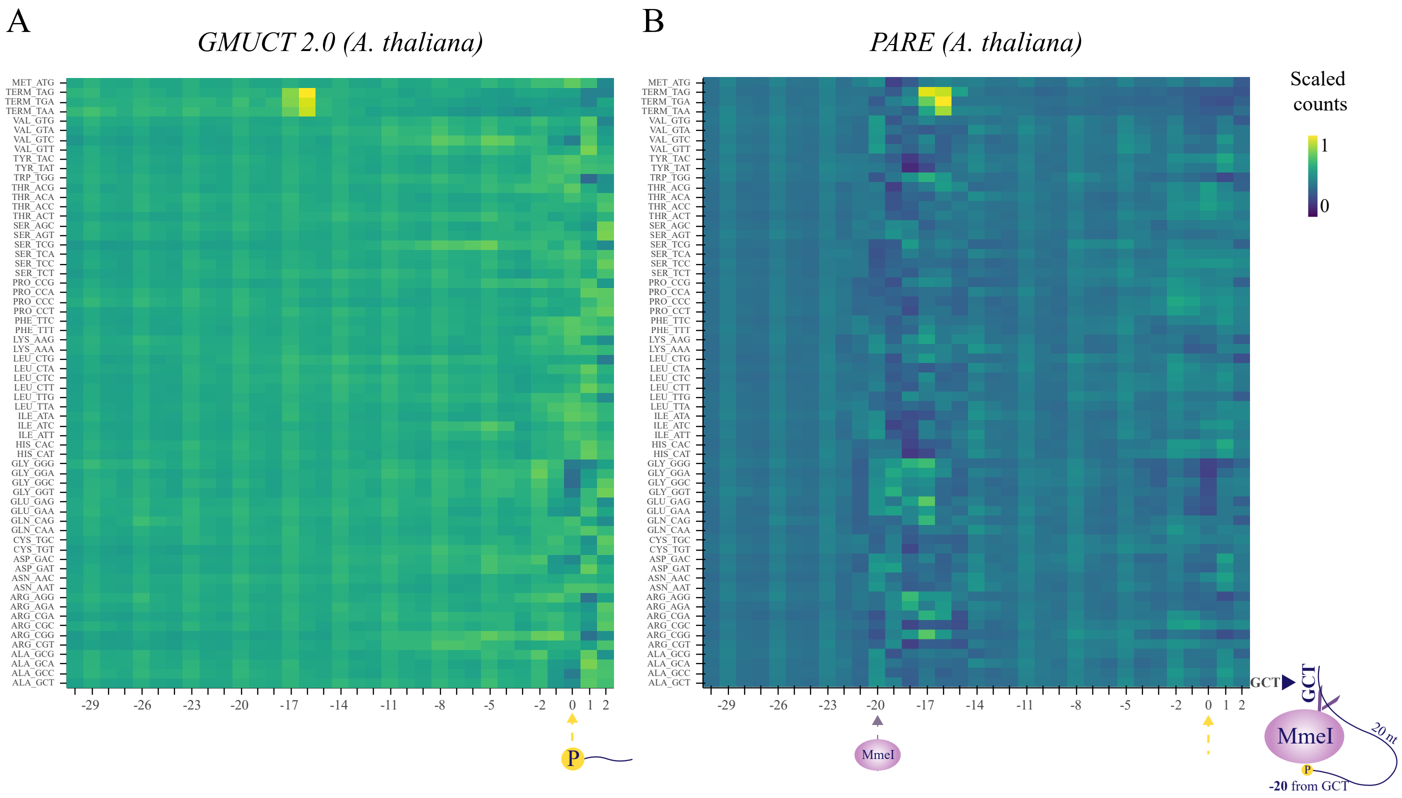


**Figure S2. Codon-specific patterns and biases in degradome sequencing libraries produced by different protocols.** Heatmaps displaying the relative 5’P degradome coverage in respect to codons at varying distances from each codon. Position 0 corresponds to the first nucleotide of the codon. Counts at this position reflect, in addition to the abundance of specific 5’P fragments, the expected single-stranded ligation bias (as all the fragments used to construct each line metagene have exactly the same sequence at positions 0,1 and 2). **A.** *A. thaliana* 5’ degradome sequencing using GMUCT 2.0 (Poethig, Meyers, Willmann and McCormick, GSE72505), where only clear ligation bias around position 0 can be observed (similar to what is expected in 5PSeq or HT-5PSeq). **B.** *A. thaliana* 5’ degradome sequencing using PARE. Aside from the 5’P ligation bias, there is also a noticeable bias observed 20nt upstream from every codon. This bias is likely introduced by MmeI, which cleaves 20nt downstream from the 5’P ends of the transcripts. Although cleavage efficiency is not supposed to be dependent on the cleavage site, it is clear that the sequence context around the cleavage site (at 0) influences the ability of MmeI to cleave the fragment and thus introduce a sequence specific bias 20 to 17 nt upstream of the cleavage site. This region overlaps with the expected ribosome protection region and this complicates codon-specific protection analysis.

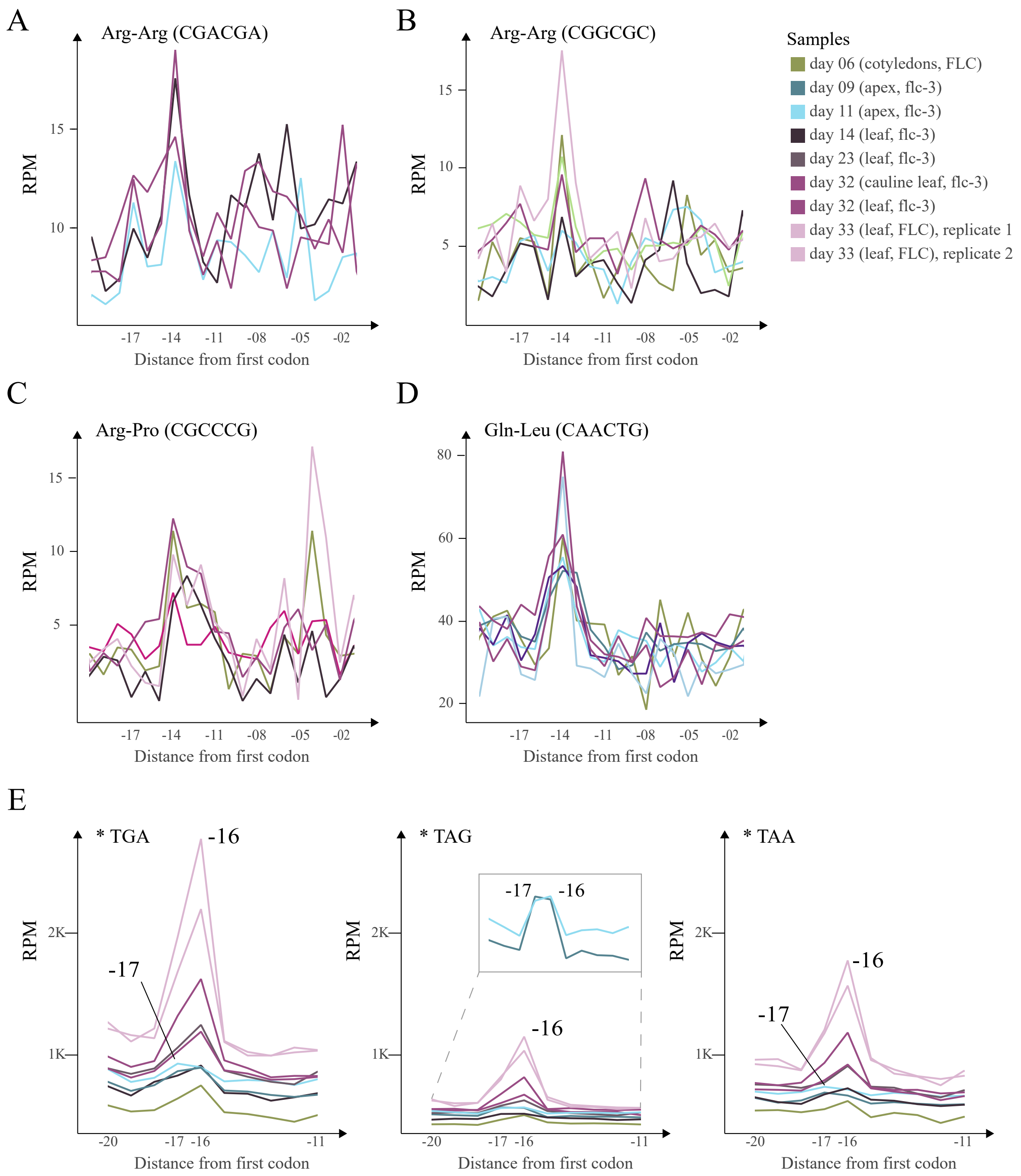


**Figure S3. Ribosome protection at commonly pausing associated codon pairs and termination codons in *A. thaliana*.** Linecharts showing metagene coverage for 5'P read in respect to selected codon features. Library size normalized counts are presented as reads per million (RPM) considering only those reads mapping in the coding regions. **A-D.** Ribosome pauses in respect to selected di-codon features **E.** *In vivo* ribosome protection for ribosomes paused at translation termination.

**Table S1.** Genes with changes in ribosome protection frame of preference between wild type and the *tif5A1-3 mutant.* Frame indicates the preferred protection frame in the indicated strain.

| Frame | Strain | Protein names | Gene name |
|---|---|---|---|
| **F0** | wild type | Protein arginine N-methyltransferase HSL7 (EC 2.1.1.320) (Histone synthetic lethal protein 7) (Type II protein arginine N-methyltransferase) (Type II PRMT) | HSL7 |
| **F0** | wild type | Protein CWH43 (Calcofluor white hypersensitive protein) | CWH43 |
| **F0** | wild type | 40S ribosomal protein S13 (S27a) (Small ribosomal subunit protein uS15) (YS15) | RPS13 |
| **F0** | wild type | DNA damage checkpoint protein LCD1 (DNA damage checkpoint protein 2) (Lethal, checkpoint-defective, DNA damage-sensitive protein 1) | LCD1 |
| **F0** | wild type | Kinetochore protein NDC80 (80 kDa spindle component protein) (Nuclear division cycle protein 80) (Two-hybrid interaction with DMC1 protein 3) | NDC80 |
| **F0** | wild type | Transcriptional activator HAC1 | HAC1 |
| **F0** | wild type | Outer spore wall assembly protein SHE10 (Sensitivity to high expression protein 10) | SHE10 |
| **F0** | wild type | Transcription factor PDR1 (Pleiotropic drug resistance protein 1) | PDR1 |
| **F0** | wild type | Protein BIR1 | BIR1 |
| **F0** | wild type | Vacuolar protein sorting-associated protein 70 (EC 3.4.-.-) | VPS70 |
| **F0** | wild type | Protein CSF1 (Cold sensitive for fermentation protein 1) | CSF1 |
| **F0** | wild type | U3 small nucleolar RNA-associated protein 12 (U3 snoRNA-associated protein 12) (DOM34-interacting protein 2) (U three protein 12) | DIP2 |
| **F0** | wild type | Putative uncharacterized protein ART2 (Antisense to ribosomal RNA transcript protein 2) | ART2 |
| **F0** | wild type | 60S ribosomal protein L31-B (L34) (Large ribosomal subunit protein eL31-B) (YL28) | RPL31B |
| **F0** | wild type | Helicase SEN1 (EC 3.6.4.-) (tRNA-splicing endonuclease positive effector) | SEN1 |
| **F0** | wild type | Protein SST2 | SST2 |
| **F0** | wild type | Probable transcription factor TDA9 (Topoisomerase I damage affected protein 9) | TDA9 |
| **F0** | wild type | Endoplasmic reticulum vesicle protein 25 | ERV25 |
| **F0** | wild type | Uncharacterized protein YMR124W | |
| **F0** | wild type | Putative uncharacterized membrane protein YMR173W-A | |
| **F0** | wild type | Ubiquitin domain-containing protein DSK2 | DSK2 |
| **F0** | wild type | Glucose-6-phosphate 1-dehydrogenase (G6PD) (EC 1.1.1.49) | ZWF1 |
| **F0** | wild type | Protein BNI4 | BNI4 |
| **F0** | wild type | Anaphase-promoting complex subunit 1 | APC1 |
| **F0** | wild type | Ribosome biogenesis protein RLP7 (Ribosomal protein L7-like) | RLP7 |
| **F0** | wild type | Restriction of telomere capping protein 1 (SEH-associated protein 2) | RTC1 |
| **F0** | wild type | Translation machinery-associated protein 46 (DRG family-regulatory protein 1) | TMA46 |
| **F0** | wild type | tRNA pseudouridine synthase 1 (EC 5.4.99.-) (tRNA pseudouridylate synthase 1) (tRNA-uridine isomerase 1) | PUS1 |
| **F0** | wild type | Putative protease AXL1 (EC 3.4.24.-) | AXL1 |

| | | | |
|---|---|---|---|
| **F0** | wild type | Protein SCD6 | SCD6 |
| **F0** | wild type | Transcription factor SPN1 (Interacts with SPT6 protein 1) (Suppresses postrecruitment functions protein 1) | SPN1 |
| **F0** | wild type | E3 ubiquitin-protein ligase linker protein MMS1 (Methyl methanesulfonate-sensitivity protein 1) (Regulator of Ty1 transposition protein 108) (Synthetically lethal with MCM10 protein 6) | MMS1 |
| **F0** | tif5A1-3 | Reduced viability upon starvation protein 161 | RVS161 |
| **F0** | tif5A1-3 | Putative uncharacterized protein YDR154C | |
| **F0** | tif5A1-3 | ATP synthase subunit 9, mitochondrial (Lipid-binding protein) (Oligomycin resistance protein 1) | OLI1 |
| **F0** | tif5A1-3 | Metallothionein expression activator | ACE2 |
| **F0** | tif5A1-3 | Tubulin alpha-3 chain | TUB3 |
| **F0** | tif5A1-3 | Altered inheritance of mitochondria protein 44 | AIM44 |
| **F2** | wild type | 40S ribosomal protein S11-B (RP41) (S18) (Small ribosomal subunit protein uS17-B) (YS12) | RPS11B |
| **F2** | wild type | Serine hydroxymethyltransferase, mitochondrial (SHMT) (EC 2.1.2.1) (Glycine hydroxymethyltransferase) (Serine methylase) | SHM1 |
| **F2** | wild type | ADP-ribosylation factor GTPase-activating protein GCS1 (ARF GAP GCS1) | GCS1 |
| **F2** | wild type | 60S ribosomal protein L4-B (L2) (Large ribosomal subunit protein uL4-B) (RP2) (YL2) | RPL4B |
| **F2** | wild type | 40S ribosomal protein S11-A (RP41) (S18) (Small ribosomal subunit protein uS17-A) (YS12) | RPS11A |
| **F2** | wild type | 60S ribosomal protein L16-A (L13a) (L21) (Large ribosomal subunit protein uL13-A) (RP22) (YL15) | RPL16A |
| **F2** | wild type | Cytochrome b mRNA maturase bI2 | BI2 |
| **F2** | wild type | Nuclear localization sequence-binding protein (p67) | NSR1 |
| **F2** | wild type | Peptidyl-prolyl cis-trans isomerase CPR6 (PPIase CPR6) (EC 5.2.1.8) (Rotamase CPR6) | CPR6 |
| **F2** | wild type | rRNA biogenesis protein RRP5 (Ribosomal RNA-processing protein 5) (U3 small nucleolar RNA-associated protein RRP5) (U3 snoRNA-associated protein RRP5) | RRP5 |
| **F2** | wild type | 40S ribosomal protein S10-A (Small ribosomal subunit protein eS10-A) | RPS10A |
| **F2** | wild type | Heat shock protein homolog SSE1 (Chaperone protein MSI3) | SSE1 |
| **F2** | wild type | GTP-binding protein RHO1 (Rho-type GTPase 1) | RHO1 |
| **F2** | tif5A1-3 | 60S ribosomal protein L2-B (L5) (Large ribosomal subunit protein uL2-B) (RP8) (YL6) | RPL2B |
| **F2** | tif5A1-3 | 40S ribosomal protein S0-A (Nucleic acid-binding protein NAB1A) (Small ribosomal subunit protein uS2-A) | RPS0A |
| **F2** | tif5A1-3 | Putative uncharacterized protein YLR154W-B | |
| **F2** | tif5A1-3 | Ferric/cupric reductase transmembrane component 1 (EC 1.16.1.9) (Ferric-chelate reductase 1) | FRE1 |
| **F2** | tif5A1-3 | 60S acidic ribosomal protein P0 (A0) (L10e) (Large ribosomal subunit protein uL10) | RPP0 |
| **F2** | tif5A1-3 | Ankyrin repeat-containing protein YAR1 | YAR1 |

**Table S2.** Candidate genes with possible ribosome frameshifts (RF) in the wild type strain of *S. cerevisiae.*

| Possible RF | Gene | Description | Similar sequences with known RF |
|---|---|---|---|
| -1 | *RPS13* | 40S ribosomal protein S13 (S27a) (Small ribosomal subunit protein uS15) (YS15) | P39470 (*Sulfolobus acidocaldarius*) |
| -1 | *RPL31B* | 60S ribosomal protein L31-B (L34) (Large ribosomal subunit protein eL31-B) (YL28) | |
| -1 | *SST2* | Protein SST2 | |
| -1 | *RLP7* | Ribosome biogenesis protein RLP7 (Ribosomal protein L7-like) | |
| -1 | *TMA46* | Translation machinery-associated protein 46 (DRG family-regulatory protein 1) | Q3TIV5 (*Mus musculus*) |
| -1 | *PUS1* | tRNA pseudouridine synthase 1 (EC 5.4.99.-) (tRNA pseudouridylate synthase 1) (tRNA-uridine isomerase 1) | Q9WU56 (*Mus musculus*) |
| -1 | *SCD6* | Protein SCD6 | |
| +1 | *RPL4B* | 60S ribosomal protein L4-B (L2) (Large ribosomal subunit protein uL4-B) (RP2) (YL2) | |
| +1 | *RPS11A* | 40S ribosomal protein S11-A (RP41) (S18) (Small ribosomal subunit protein uS17-A) (YS12) | P0CT73 (*Schizosaccharomyces pombe*) |
| +1 | *RPL16A* | 60S ribosomal protein L16-A (L13a) (L21) (Large ribosomal subunit protein uL13-A) (RP22) (YL15) | |
| +1 | *CPR6* | Peptidyl-prolyl cis-trans isomerase CPR6 (PPIase CPR6) (EC 5.2.1.8) (Rotamase CPR6) | |
| +1 | *RHO1* | GTP-binding protein RHO1 (Rho-type GTPase 1) | |
| +1 | *RPS10A* | 40S ribosomal protein S10-A (Small ribosomal subunit protein eS10-A) | |
